# Development of a fluorescent quantitative real-time polymerase chain reaction assay for the detection of novel-goose parvovirus in vivo

**DOI:** 10.1101/2021.07.09.451858

**Authors:** Haibin Ma, Yahui Li, Junzheng Yang

**Author notes:** Correspondence to Junzheng Yang, Bioland Laboratory (Guangzhou Regenerative Medicine and Health Guangdong Laboratory), Guangzhou, 510700, China.

## Abstract

**Objectives:** To develop a sensitive, highly specific fluorescent quantitative real-time PCR assay for accurate detection and quantification of novel-goose parvovirus (N-GPV) in vitro and in vivo.

**Methods:** Specific primers was designed based on N-GPV inverted terminal repeats region; virus RNA (DFV, NDV, AIV, DHV-1, DHV-3) and virus DNA (MDPV, GPV, N-GPV) were extracted, cDNA (DFV, NDV, AIV, DHV-1, DHV-3) were prepared from viral RNAs using M-MLV Reverse Transcriptase, and prepared cDNA (DFV, NDV, AIV, DHV-1, DHV-3) and DNA (MDPV, GPV, N-GPV) amplified by real-time PCR; the sensitivity, specificity and reproducibility of established real-time PCR methods were evaluated, and finally we validated the reliability of real-time PCR methods in ducklings models in vivo.

**Results:** The standard curve of established real-time PCR had a good linearity (slope was −0.3098, Y-intercept was 37.865, efficiency of standard curve was 0.995); the detection limit of established real-time PCR for N-GPV was 10 copies/reaction. The sensitivity of real-time PCR was 10 copies/μL, which was 1000 times higher than conventional gel-based PCR assay. The results of intra-assay CVs (0.04-0.74%) and inter-assay CVs (0.16-0.53%) showed that the real-time PCR assay had an excellent repeatability. This method also could efficiently detect viral load in heart, liver, spleen, lung, kidney, pancreas, bursa of Fabricius, brain, blood and excrement from ducklings models after N-GPV infection from 6h to 28 days, which could provided us a dynamic distribution observation of N-GPV viral load using this real-time PCR assay in vivo.

**Conclusion:** In the study, we developed a high sensitive, specific and reproducible real-time PCR assay for N-GPV detection. The established real-time PCR assay was suitable for parvovirus detection and quantification simultaneously, no matter sample obtained from blood, internal organs or ileac contents; the present work may provide insight into the pathogenesis of N-GPV and will contributes to better understanding of this newly emerged novel GPV related virus in cherry valley ducks.

## 1 Introduction

There are several parvovirus duck strain variations on the basis of phylogeny and homology analysis main including GPV (goose parvovirus), MDPV (muscovy duck parvovirus) and N-GPV (novel-goose parvovirus). The structure of GPV and MDPV contain a linear, single-stranded DNA genome (5.0-5.1kb) having two inverted terminal repeats (ITR) at both ends forming a hairpin structure containing stem and bubble region having length 456 and 442 nucleotides for MDPV and GPV respectively^[1, 2]^. They have two open reading frames (ORFs), left and right ORFs encode non-structural (NS1 and NS2) and structural (VP1, VP2 and VP3) proteins respectively^[3, 4]^.

N-GPV is a kind of newly emerged, distinct GPV-related virus usually causes short beak dwarfism syndrome (SBDS) in young ducks. In China, it was reported initially in 2014 and since then it severely affected the duck industry mainly due to reduction in weight of animals and size of beak, there are numerous SBDS outbreaks in mule duck and Cherry Valley duck flocks in various regions of China^[5, 14]^. GPV is an etiological agent for Derzsy’s disease, mortality rate is 70-100% during first four weeks, usually resulted in huge economic losses to the duck industry^[5, 6]^. Clinically, the disease is characterized by lethargy, stunting, anorexia, locomotor dysfunction, watery diarrhea and death within 3-5 days^[7]^. MDPV characterized clinically by signs almost similar to GPV mainly affects three-week-old Muscovy ducklings^[1, 8, 9]^. The complete genome sequence of both parvoviruses shares 79.7-85.0% homology^[10, 11,18]^.

SBDS caused by novel GPV-related virus (N-GPV) was first reported in France during 1970s in mule ducks^[12, 13]^. Adult ducks are found resistant to this disease while young animals infected with N-GPV have swollen tongue, shorter tibia and stunted growth with 2% mortality rate while morbidity reaches 10%-30% in most cases^[15]^. The losses to the duck industry therefore are mainly due to reduction in weight and size of animals^[16]^.

Real-time PCR is a new technique with remarkable sensitivity, specificity, accuracy, reproducibility, visualization of results, simple and less-contamination potential compared to other diagnostic methods, it has become a potentially powerful alternative in microbiological diagnostics ^[17]^. In order to effectively prevent SBDS in duck and reduce the economic loss caused by SBDS, combining the advantages of real time PCR, in this article, we developed a fully validated, highly specific and reliable real-time PCR assay for detection of N-GPV infection, and validated effectiveness and specificity of real-time PCR assay in vitro and in vivo.

## 2. Methods

### 2.1 Preparation of Virus and PCR template DNA

N-GPV AH strain was isolated from Cherry Valley duck flock from Anhui province in 2021 in our lab. GPV and MDPV were bought from China Veterinary Culture Collection Center. Duck flavivirus (DFV), Newcastle disease virus (NDV), Avian influenza (AIV), Duck hepatitis virus type 1 and type 3 (DHV-1 and DHV-3), *Riemerella anatipestifer* (*R. a.*), *Escherichia coli* (*E. coli*.) were stored in our lab.

Viral RNA (DFV, NDV, AIV, DHV-1, DHV-3) and DNA (MDPV, GPV, N-GPV) were extracted using Viral DNA/RNA Miniprep Kit (Axygen, New York, USA), according to the manufacturer’s instructions. cDNA (DFV, NDV, AIV, DHV-1, DHV-3) were prepared with viral RNAs using M-MLV Reverse Transcriptase (Promega, Wisconsin, USA). Bacterial genomic DNA (*R. a*. and *E. coli*) were extracted using TIANamp Bacteria DNA Kit (Tiangen, Beijing, China). All DNA and cDNA were stored at −80°C until for use.

### 2.2 PCR primer design

The primers of real-time PCR assay were designed using Integrated DNA Technologies (Primer software, Skokie, Illinois, USA) on the basis of highly conserved regions of N-GPV. The conserved regions having characteristic variation with GPV and MDPV were selected and the primer sequence were shown at Table 1. Primers were verified by Basic Local Alignment Search Tool for specificity analysis and synthesized by Thermo Scientific Co., Ltd. (Guangzhou, China).

**Table 1:**
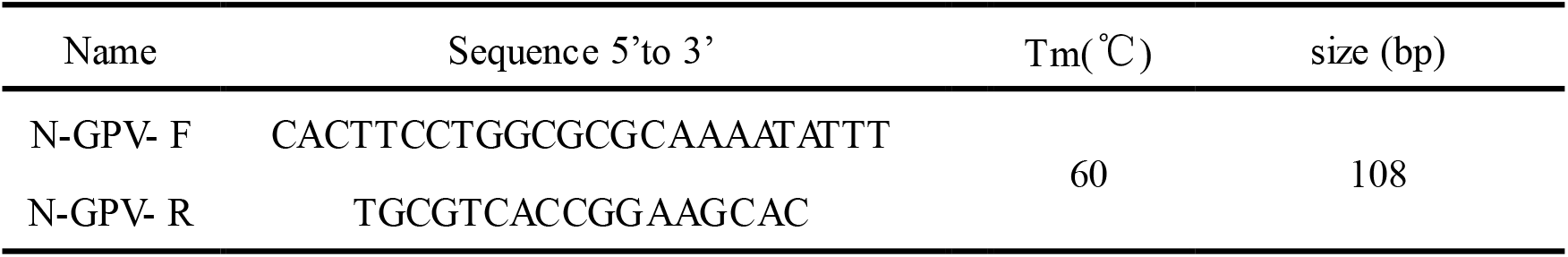
Sequences of the primers used in real-time PCR method for N-GPV detection

### 2.3 Preparation of standard plasmid DNA templates

The DNA templates length of N-GPV (strain AH, GenBank Accession No. MH444513) was 4,639-4,748bp amplified by real-time PCR and conventional PCR. The product was ligated into the pMD-18T vector (TaKaRa, Dalian, China) and transformed into *E.coli* DH5α competent cells (Tian, Beijing, China). The pMD-18T-ITR was extracted using Plasmid Extraction Minin Kit (Solarbio, Beijing, China) and was used as the standard plasmid. The recombinant plasmid was quantified using a NANO-DROP 2000 spectrophotometer (Thermo Scientific, Waltham, MA, USA). The plasmid copy number was calculated using the equations described by Yun et. al. on the basis of the molecular weight^[19, 20]^.

### 2.4 Real-time PCR assay

The real-time PCR was performed using the Bio-RAD/CFX384 Real-time PCR system (Bio-Rad, Hercules, CA) according to the manufacturer’s instructions. The optimized PCR reaction system were as follows: 2μL DNA template, 10μL Go-tag qPCR Master Mix (Promega, Wisconsin, USA), 0.2μL of forward primer, 0.2μL of reverse primer (N-GPV-F and N-GPV-R, 20μmol/L), and 7.6μL autoclaved nanopure water. Each reaction cycles comprised an initial activation step at 95 °C for 2 min, followed by 40 cycles at 95 °C for 15 sec, 60 °C for 1 min. The fluorescence signal was collected during annealing/extension step. Ten-fold serial dilution concentrations of recombinant plasmid pMD-18T-ITR containing different copy numbers (1×10 ^9^ to 1×10 DNA copies/μL) were used to generate the standard curve. Analysis of each assay was carried out using Bio-Rad CFX Manager (Bio-Rad, Hercules, CA) according to manual instructions.

### 2.5 Sensitivity analysis

The sensitivity analysis of real-time PCR and conventional PCR was determined by using different recombinant plasmid pMD-18T-ITR concentrations. pMD-18T-ITR template was prepared as follows: 10-fold serial dilution concentrations ranging from 1×10 ^9^copies/μL to 1×10copies/μL, 2μL of each dilution sample was used as a template for specificity analysis, and each concentration of sample repeated two times, PCR products were visualized in 1.0% agarose gel provided by the manufacturer.

### 2.6 Specificity and reproducibility analysis

DNA from MDPV, GPV, N-GPV and other pathogens including DFV, NDV, AIV, DHV-1, DHV-3, *R. a.* and *E. coli.* was used to validate the specificity of the established real-time PCR assay for N-GPV detection. The reproducibility of real-time PCR assay was accessed by analyzing mean coefficient of variation (CV) by using serial dilution concentrations of pMD-18T-ITR (10^7^copies/ μL, 10^6^copies/μL, 10^5^copies/μL, 10^4^copies/μL). The intra-assay variability of real-time PCR assay was accessed by using each dilution concentration in triplicate and comparing the coefficient of variation. The coefficient of variation expresses the standardized measure of dispersion with time for inter-assay variability.

### 2.7 Animal experiments

All procedures involving animal experiment were approved by the Institute of Animal Husbandry and Veterinary, South China Agricultural University, China. Sixty 1-day old Cherry Valley ducklings obtained from hatchery (Guiliu Fowl Company, Guangdong, China) were kept at experimental station under controlled conditions for acclimatization to the new environment. The ducklings were then randomly divided into treatment group and control group (30 ducklings/group), housed separately and bred in a parvovirus-free environment. All the ducklings were free of parvovirus-specific maternal antibodies confirmed by result of ELISA detection. For comparison, the ducklings in treatment group were injected intramuscularly with 500 μL of AH or GD viral allantoic fluid and the control group injected with 500 μL PBS. The ducklings in treatment group and control group were kept for 4 weeks and all humane procedures and biosecurity guidelines were followed during experiment.

Three ducks from control group and Three ducks from treatment group were randomly selected and killed at each time point. Heart, liver, spleen, lung, kidney, pancreas, bursa of Fabricius, brain, excrement and blood from the ducks in control group and treatment group were analyzed by real-time PCR at 6 h, 12 h, day 1, day 3, day 7, day 14, day 21 and day 28 after virus infection.

In detail, obtained samples were homogenized in PBS (20%, w/v), viral DNA were extracted from tissue homogenates using Viral DNA/RNA Miniprep Kit (Axygen, New York, USA). The viral load of each sample was quantified using real-time PCR assay.

## 3 Results

### 3.1 Establishment of real-time PCR standard curve

For establishment of real-time PCR standard curve, the concentration of standard pMD-18T-ITR plasmid DNA was quantified (344ng/μL) and OD_260_/OD_280_ of standard pMD-18T-ITR plasmid DNA was 1.90; the copy numbers of pMD-18T-ITR plasmid DNA were 8.7×10 ^10^copies/ μL. The real-time PCR amplification curves were generated by pMD-18T-ITR plasmid DNA at the concentration 1×10 ^7^copies/μL, 1×10 ^6^copies/μL, 1×10 ^5^copies/μL, 1×10 ^4^copies/μL, and 1×10 ^3^copies/ μL. The generated standard curve showed linearity with a slope of −0.3098 and the Y-intercept was 37.865, efficiency of standard curve was 0.995(conducted by software of Bio-Rad CFX Manager (Bio-Rad, Hercules, CA) ) (Figure 1).

**Figure 1.**
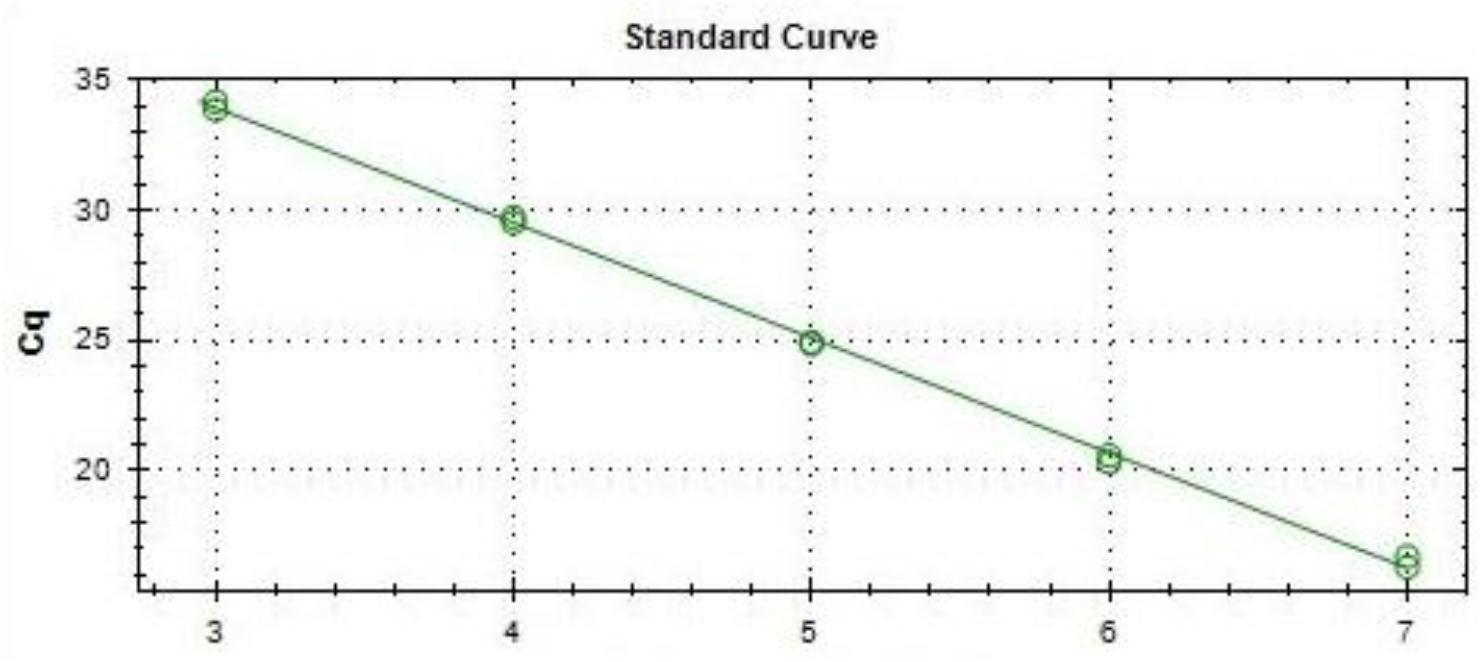
Establish the standard curve of pMD-18T-ITR plasmid DNA by real-time PCR. Serial dilution concentrations of standard DNA ranging from 10^7^-10^3^ (copies/μL) were used. The slope of standard curve was −0.3098, the Y-intercept was 37.865, and efficiency of standard curve was 0.995.

### 3.2 Sensitivity, specificity, reproducibility and dynamic range analysis of established real-time PCR

To evaluate the sensitivity of real-time PCR assay, serial dilution concentrations of pMD-18T-ITR plasmid DNA were used. The results showed that the detection limit of real-time PCR was 10 copies/reaction, the detection limit of conventional PCR was 1×10 ^4^ copies/reaction; sensitivity of real-time PCR method was 1000-times higher than conventional PCR (10copies/reaction versus 1×10 ^4^copies/reaction) (Figure 2 and Figure 3).

**Figure 2.**
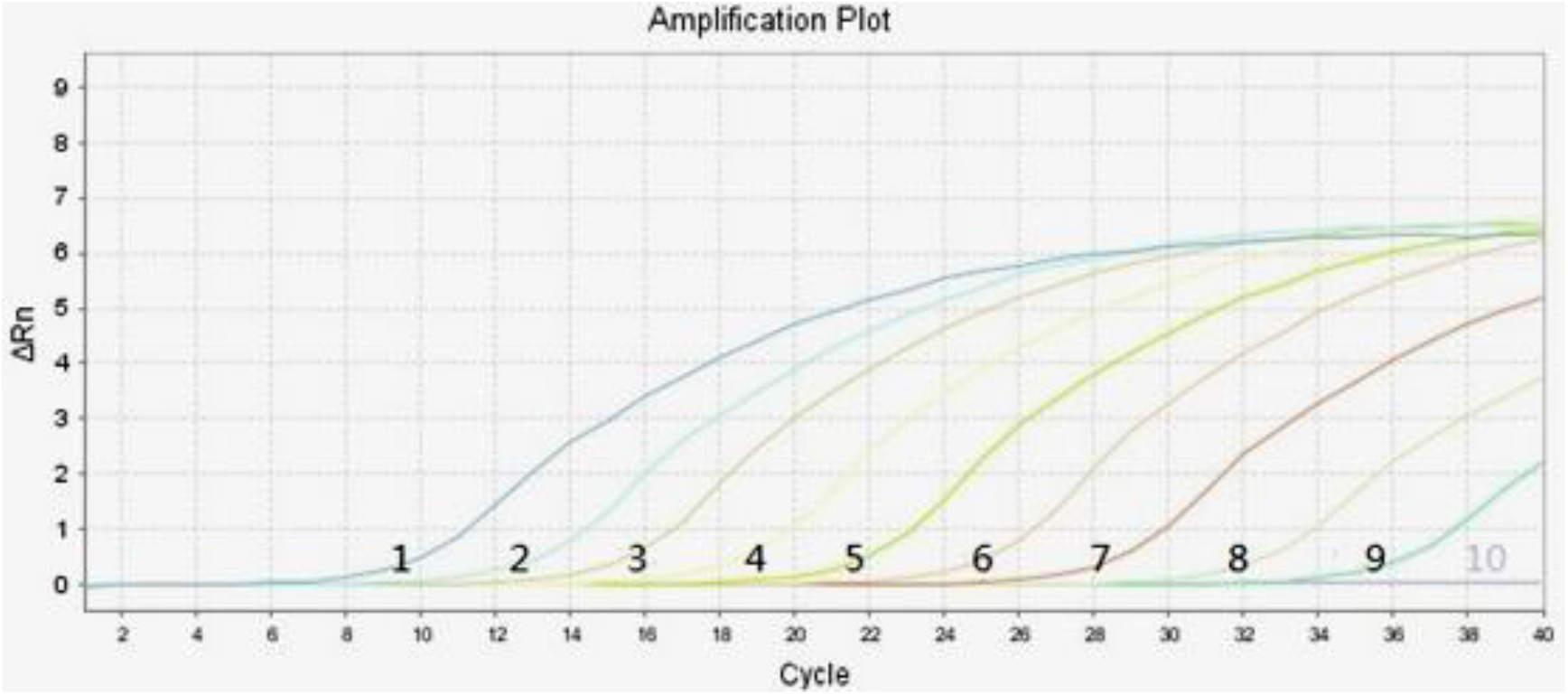
Amplification results of pMD-18T-ITR plasmid DNA by established real-time PCR for sensitivity verification. Ten-fold serial dilution concentrations of the DNA standard sample ranging from 1×10 ^9^copies/μL to 1 ×10 ^1^copies/μL were used. Number 1-9 indicated different DNA sample concentration in turn: 1×10 ^9^copies/μL, 1×10 ^8^copies/μL, 1×10 ^7^copies/μL, 1×10 ^6^copies/μL, 1×10 ^5^copies/μL, 1×10 ^4^copies/μL, 1×10 ^3^copies/μL, 1×10 ^2^ copies/μL, 1×10 ^1^ copies/μL; number 10: blank control).

**Figure 3.**
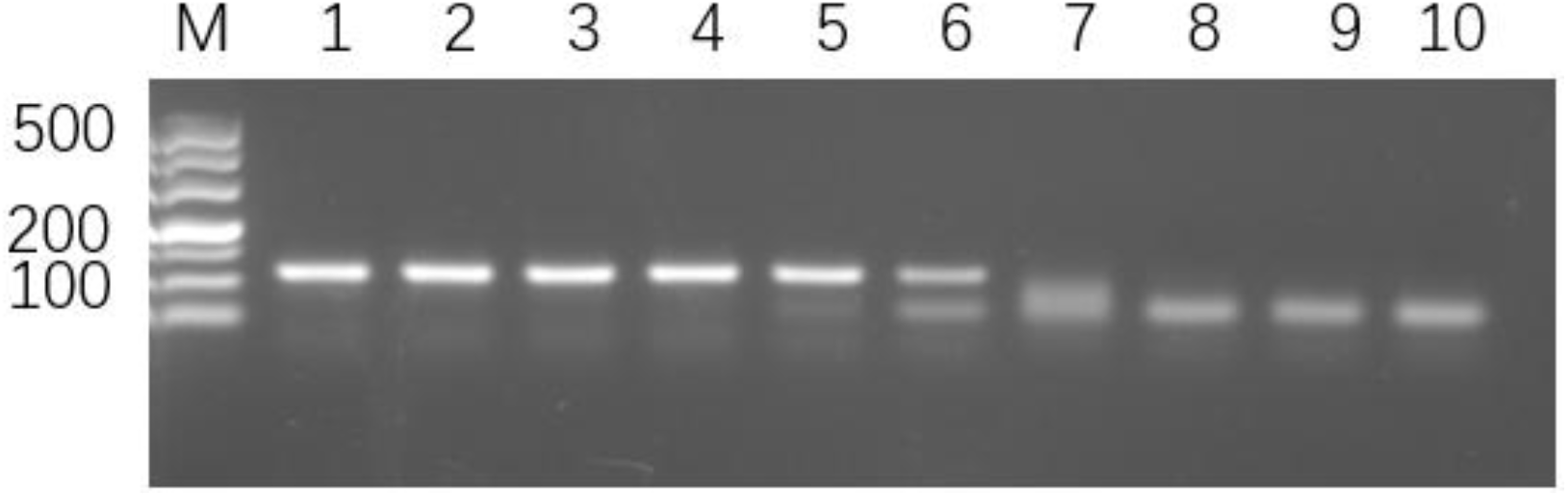
Results of conventional PCR for pMD-18T-ITR plasmid DNA detection for sensitivity verification. Ten-fold serial dilution concentrations of DNA standard samples ranging from 1×10 ^9^copies/μL to 1×10copies/μL were detected by conventional PCR. Number 1-9 indicated different DNA sample concentration in turn: 1×10 ^9^copies/μL, 1×10^8^copies/μL, 1×10^7^copies/μL, 1×10 ^6^copies/μL, 1×10 ^5^copies/μL, 1×10 ^4^copies/μL, 1×10 ^3^copies/μL, 1×10 ^2^copies/μL, 1×10 ^1^ copies/μL; number 10: blank control.

To evaluate the specificity of real-time PCR assay, virus DNA (MDPV, GPV, N-GPV), virus cDNA (DFV, NDV, AIV, DHV-1, DHV-3), bacterial DNA (*R. a., E. coli*) and control (nuclease-free water) were used, the results demonstrated that there was a smooth amplification curve for N-GPV DNA, there were no amplification curve for virus DNA (MDPV, GPV), virus cDNA (DFV, NDV, AIV, DHV-1, DHV-3), bacterial DNA (*R. a., E. coli*) and control (nuclease-free water) (Figure 4).

**Figure 4.**
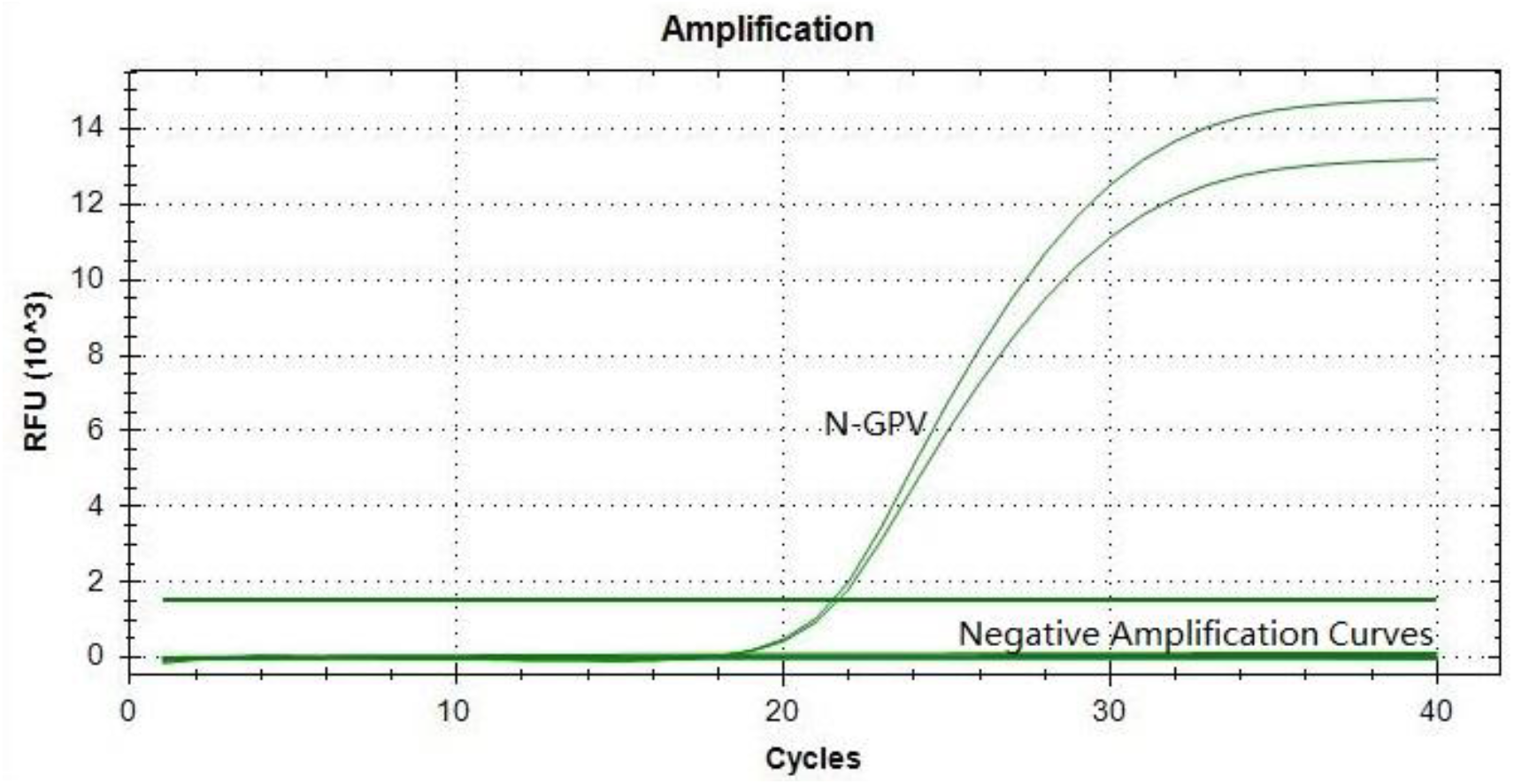
Specificity verification of the established real-time PCR. Nine DNA or cDNA samples from different viruses and bacteria (MDPV, GPV, DFV, NDV, AI V, DHV-1, DHV-3, *R. a.*, *E. coli* ) were used to evaluate specificity of established real-time PCR assay.

For validation of repeatability of established real-time PCR assay, four different concentrations of standard DNA (1×10 ^7^copies/ μL, 1×10 ^6^copies/μL, 1×10 ^5^copies/μL, 1×10 ^4^copies/μL) were used, and every concentration of standard DNA repeated three times for statistics analysis. The results showed that The intra-assay CVs were in the range of 0.04-0.74%, the inter-assay CVs were in the range of 0.16-0.53% (Table 2). These demonstrated that the established real-time PCR assay had a high reproducibility for N-GPV DNA detection.

**Table 2.** Reproducibility validation of established real-time PCR

| Samples | Copies | n | Intra-batch (Ct Value) |  |  | Inter-batch (Ct Value) |  |  |
| --- | --- | --- | --- | --- | --- | --- | --- | --- |
|  |  |  | Average | SD | CV(%) | Average | SD | CV(%) |
| 1 | $10^7$ | 3 | 19.61 | 0.15 | 0.74 | 30.97 | 0.10 | 0.53 |
| 2 | $10^6$ | 3 | 23.05 | 0.01 | 0.04 | 27.67 | 0.04 | 0.16 |
| 3 | $10^5$ | 3 | 27.70 | 0.11 | 0.39 | 23.02 | 0.15 | 0.54 |
| 4 | $10^4$ | 3 | 31.00 | 0.11 | 0.34 | 19.58 | 0.07 | 0.24 |

### 3.3 Dynamic distribution of N-GPV viral load in vivo using real-time PCR assay

The reproductive properties of N-GPV AH in Cherry Valley ducks were analyzed by detection of viral load in blood, internal organs and ileac contents of infected ducks. The results showed that all tissues and blood isolated from infected ducks had high levels of viral load from 6 h to 28 days after quantification of N-GPV by using the established real-time PCR assay. The viral load of N-GPV presented a typical S-shaped curve accompanied with the passage of time, the highest viral load usually reached at day 7 or day14 and the lowest viral load usually reached at day 1 or day 3 in all tissues and blood isolated from infected ducks.

In detail, the average value of N-GPV AH were 10^5.76^ in heart, 10^5.08^ in blood, 103.35 in pancreas, 10^5.34^ in kidney, 10^4.78^ in spleen, 10^4.10^ in brain, 10^5.64^ in faeces, 104.94 in liver, 10^5.03^ in lung, and 10^5.07^ in bursa of fabricius after infection for 6h. The lowest point of viral load in these samples usually appears at day 1 (pancreas, spleen, liver, bursa of fabricius) or day 3 (heart, blood, kidney, brain, faeces, lung) and the average value was 10^3.85^ in heart), 10^4.22^ in blood, 10^2.63^ in pancreas, 10^3.45^ in kidney, 10^2.96^ in spleen, 10^3.32^ in brain, 10^4.26^ in faeces, 10^3.64^ in liver, 10^4.06^ in lung, 102.75 in bursa of fabricius. The highest point of viral load appears at day 7 (blood, kidney, spleen, brain, liver, lung, bursa of fabricius), day 14 (faeces), day 21 (heart, pancreas) and the average value was 10^7.43^ in heart, 10^8.01^ in blood, 10^6.05^ in pancreas, 107.22 in kidney, 10^8.11^ in spleen, 10^5.91^ in brain, 10^6.43^ in faeces, 10^7.19^ in liver, 10^7.07^ in lung, 10^5.58^ in bursa of fabricius (Figure 5). Those data demonstrated that real-time PCR assay was not only a powerful method for detection of the N-GPV AH in all kinds of tissues, excrement and blood from infected ducks, but also provided us a dynamic observation of the distribution of the N-GPV AH in all kinds of tissues in the body, which was of great significance in clinical diagnosis and clinical treatment.

**Figure 5.**
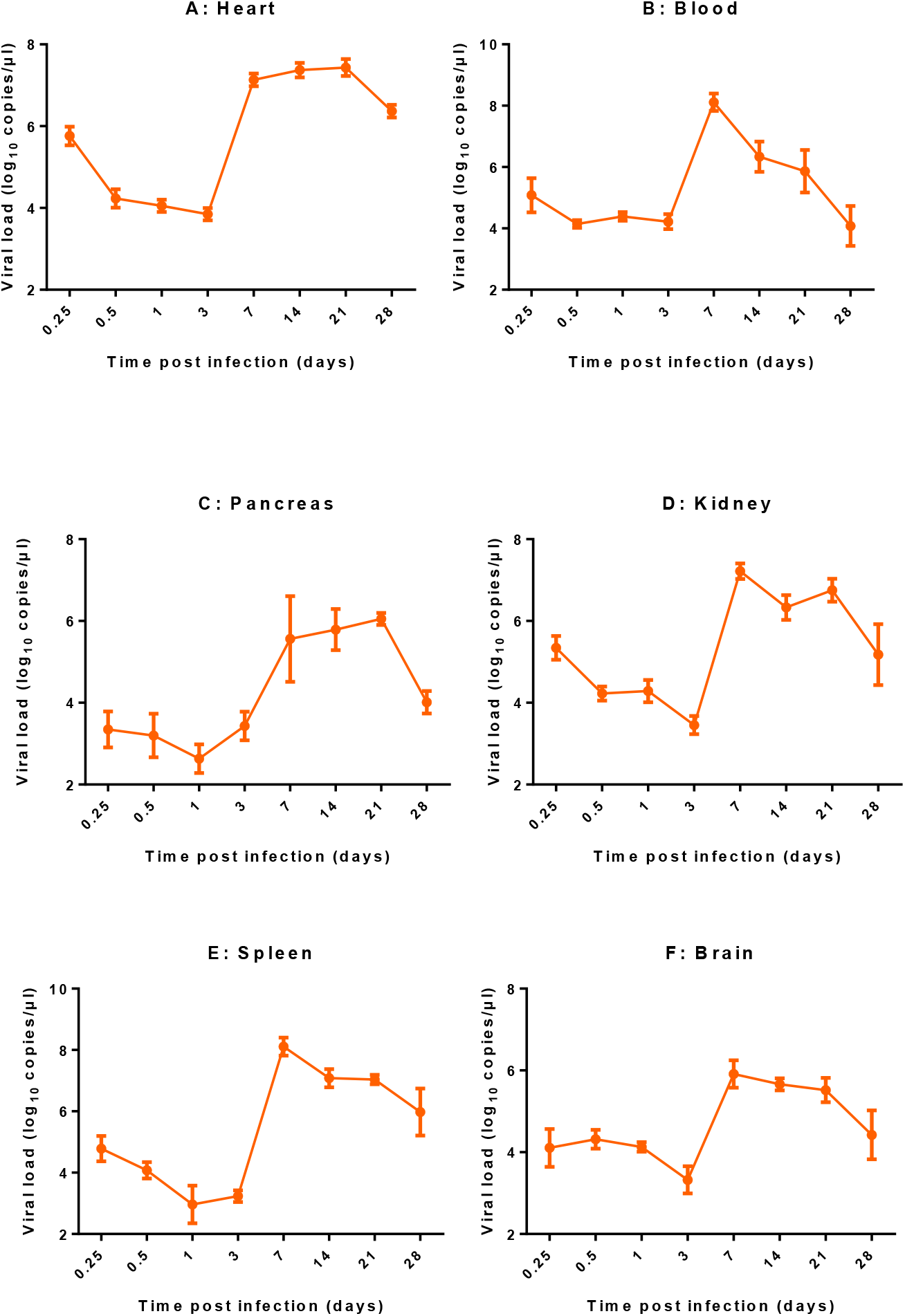
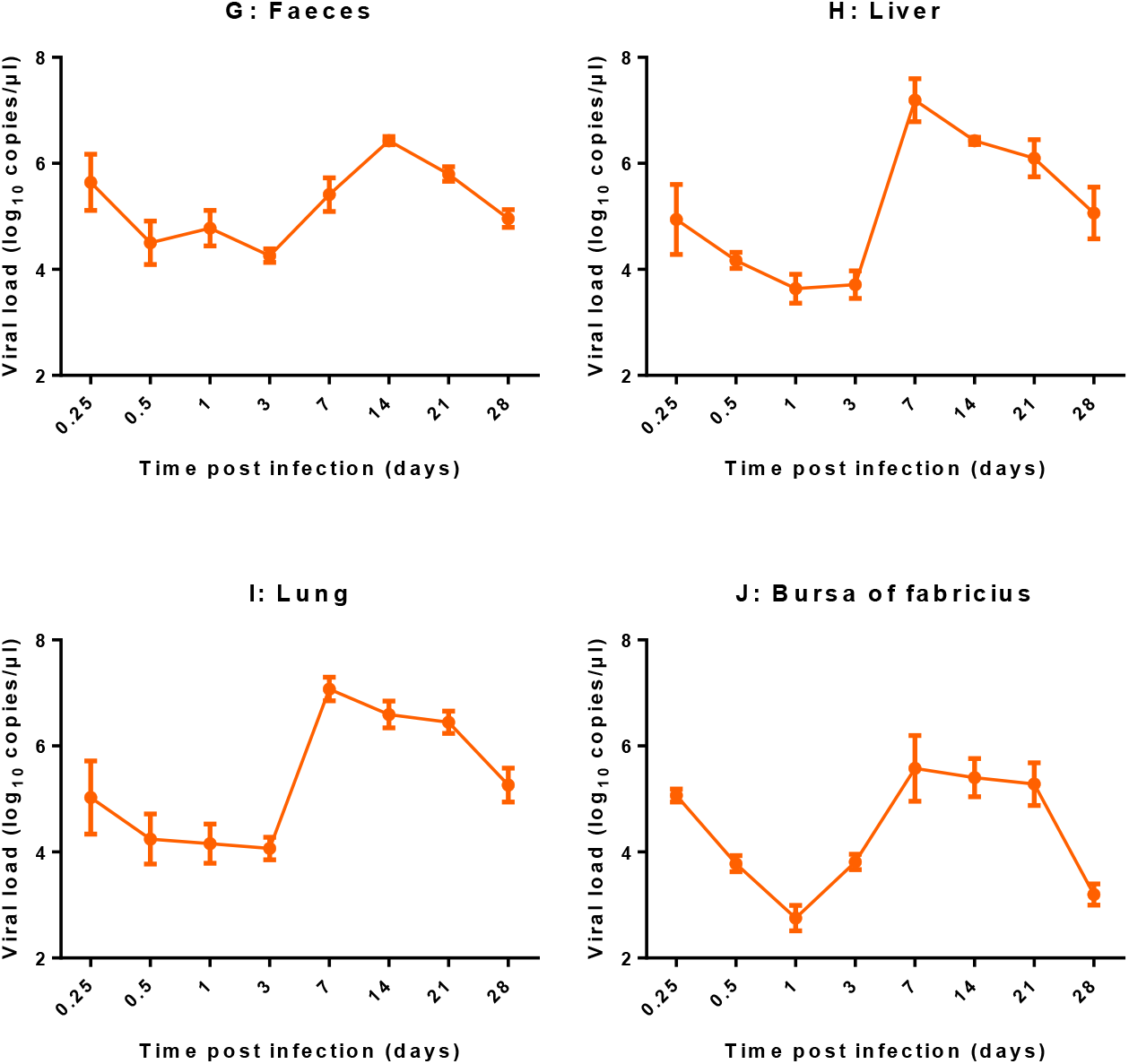
Detection results of viral loads in tissues (heart, liver, spleen, lung, kidney, pancreas, bursa of fabricius, brain), excrement and blood isolated from Cherry Valley ducks by the established real-time PCR assay at 6 h, 12 h, day 1, day 3, day 7, day 14, day 21 and day 28 after N-GPV AH infection. There were eight time points from 6h to 28 days, the result of each time point were in triplicate, the data of viral loads showed as mean±SEM.

## 4 Discussion

SBDS in ducks was first reported in France in 1970s^[13]^, followed by outbreaks in Taiwan (1989), Poland (1995) ^[21]^, Hungary (2009) and mainland China (2014)^[22, 23]^. SBDS in ducks usually caused by GPV or MDPV, and co-infection with N-GPV and MDPV strains also existed; there were several variations among these parvovirus duck strains, the sequence analysis indicated that those N-GPV variations had higher homology among them compared with classical GPV and MDPV on the basis of phylogeny and homology analysis ^[24]^.

There are many methods for the detection of GPV, N-GPV, MDPV, but those detection methods have their disadvantages. For example, virus isolation, electron microscopy, immunological-based assays which turned out to be laborious and time-consuming; conventional PCR assay and sequencing cannot effectively distinguish the various parvovirus strains because of highly homology at molecular or genomic level between GPV and N-GPV strains; Enzyme-linked immunoabsorbent assay (ELISA) methods based on VP3 protein are also applied in the detection of GPV and MDPV with a lower detection sensitivity^[25, 26]^; Loop-mediated isothermal amplification (LAMP) assay is also developed for rapid, accurate, sensitive and convenient detection of MDPV and GPV, but this method always obtains high number of false positive results^[27]^.

Here, we developed a real-time PCR assay for the quantification of N-GPV genome copies in ducks and found that this method is highly sensitive, specific and reproducible compared with conventional PCR. Moreover, the real-time PCR assay can be used for simultaneous detection and quantification of DNA. It allowed us to study the pathogenesis of disease and mechanisms of virus transmission by investigation of viral dynamics. The assay can also be used for quantification of viral load in different tissues at various times after infection; the data of viral load could be useful for multi-aspect understanding of viruses. Compared with other studies, this research reported the distribution and quantity of N-GPV and its characteristic lesions in infected muscovy ducks, which was of great significance in clinical diagnosis and clinical treatment.

The viral load of N-GPV presented a typical S-shaped curve accompanied with the passage of time, the highest viral load usually reached at day 7 or day 14 and the lowest viral load usually reached at day 1 or day 3 in all tissues and blood isolated from infected ducks. It may be because after entering the body, the virus needs an adaptation process to overcome the resistance against disease due to body’s immune system.

In summary, we had established a sensitive, specific and reproducible real-time PCR assay for N-GPV detection, it could be suitable for parvovirus detection and quantification simultaneously, and we could observe and analyze the dynamic distribution of N-GPV viral load by established real-time PCR assay in vivo, which was of great significance in clinical diagnosis and clinical treatment.

## 6 Acknowledgements

None.

